# Neuronal Activity Alters Neuron to OPC Synapses

**DOI:** 10.1101/2022.07.25.501254

**Authors:** Moura, Parvathaneni, Sahagun, Noguchi, Brennan, Tilton, Brock, Halladay, Pleasure, Cocas

## Abstract

The mechanisms that drive the timing and specificity of oligodendrocyte myelination during development, or remyelination after injury or immune attack are not well understood. Recent work has shown that oligodendrocyte progenitors receive synapses from neurons, providing a potential mechanism for neuronal-glial communication. We hypothesize that these connections are important both for correct myelination of neurons during development and for myelination during neuronal plasticity. We utilized chemogenetic tools and viral monosynaptic circuit tracing to analyze these neuroglial connections and to examine OPC proliferation, myelination, synapse formation, and neuronal-glial connectivity after increasing or decreasing neuronal activity in vivo. We found that increasing neuronal activity increased OPC activation, but not proliferation. We also found that altering neuronal activity altered neuronal-glial synaptic connections: while it did not impact the total number of neuronal inputs, or the number of inhibitory neuronal inputs, it did alter the number of excitatory neuron to OPC connections. We also found that increasing or decreasing neuronal activity impacted the ratio of excitatory and inhibitory synapses. Our data show that neuronal activity affects OPC activation, neuronal synapse formation onto OPCs, as well as the types of neuronal inputs to OPCs, indicating that neuronal activity is important for OPC circuit composition and function.

## Introduction

Fast and efficient conduction of electrical nerve impulses happens via saltatory conduction thanks to the insulation of the axons through the myelin sheath. (STAMPFLI, 1954) Myelination is necessary for the maturation of the neural circuitry required for cognition, complex motor behaviors, and sensory integration, and it also regulates the timing of activity in neural circuits and is important for maintaining the health of axons and providing nutritional support. (Saab & Nave, 2017) Oligodendrocytes, the cells responsible for producing myelin, continue to proliferate and add new myelin throughout the lifespan. Developmental myelination is complete in humans by age 40, and a secondary phase, called adaptive myelination, continues throughout life characterized by changes in myelination after motor learning, reduction of myelination with social isolation and monocular deprivation, and recovery from injury (Bechler et al., 2018) (McKenzie et al., 2014) (Liu et al., 2016) (Etxeberria et al., 2016a) (Duncan et al., 2018).

OPCs cells express a chondroitin sulfate proteoglycan and are the oligodendrocyte progenitor cells (OPCs), one of the largest pools of dividing cells in the postnatal brain. OPCs produce new oligodendrocytes that remyelinate axons after injury and in demyelinating diseases or injury such as MS and hypoxia (Wang et al., 2018). Recent work has shown that OPCs receive synapses from neurons in both white and gray matter, and are activated by the neurotransmitters glutamate and GABA from neuronal presynaptic vesicles (Bergles et al., 2000) (Lin & Bergles, 2004). Furthermore, OPCs express ion channels, including Na+, K+, and Ca++ voltage gated channels, as well as AMPA, NMDA, GABAA, and GABAB receptors (Moura et al., 2022) (Kukley et al., 2010) (Bergles et al., 2000) (Paoletti et al., 2013) (Serrano-Regal et al., 2020). The expression of these channels and receptors are developmentally regulated, peaking early in postnatal development at a time window consistent with developmental myelination. (Spitzer et al., 2019) These data suggest that neuronal-glial synapse formation is important for myelination. OPCs receive synapses from neurons: these has been shown at the level of the individual synapse via paired electrophysiological recordings in hippocampal slices to determine the functional nature of these synapses, as well as with electron microscopy to confirm the presence of presynaptic densities adjacent to OPC structures that have many of the features of a postsynaptic structure (Bergles et al., 2000)(Lin & Bergles, 2004). It has also been shown, via electrophysiological recordings, that OPCs receive both excitatory and inhibitory inputs, and that the neuronal inputs evoke small postsynaptic currents (EPSCs or IPSCs, respectively) in OPCs in the hippocampus (Bergles et al., 2000). However, little functional information on the circuit cytoarchitecture has been examined in different forebrain structures. One recent paper found neuron to OPC connections in multiple regions of the cortex but found no changes in connectivity after alterations of sensory inputs via whisker trimming. (Monje & Káradóttir, 2020)

Several lines of evidence support the idea that neuronal activity influences OPC development and myelination (Bergles et al., 2010)(Gallo et al., 1996), (Yuan et al., 1998), (Gudz et al., 2006). Increasing neuronal activity using chemogenetics leads to an increase in OPC proliferation and differentiation of OPCs, as well as increasing the probability of an axon being myelinated, and results in thicker myelin on stimulated axons. (Mitew et al., 2018a) The authors also found that decreasing neuronal activity could decrease the probability of myelination, indicating that myelination is activity-dependent and that neuronal activity is instructive for myelination. This is developmentally regulated: juvenile mice have a more robust increase in OPC proliferation after increases in neuronal activity compared to adult mice (McKenzie et al., 2014), (Gibson et al., 2014), (Mitew et al., 2018a) Blocking sensory inputs from neurons in the retina during the critical period of visual sensory input results in a decrease in myelination of the optic tract (Etxeberria et al., 2016b). Due to these earlier findings we chose to focus on juvenile mice, when the initial period of myelination is still occurring. We sought to determine whether neuronal activity directly changes OPC activity, if neuronal activity modulation altered neuron to OPC connectivity, and finally, the role of neuronal activity on synapse formation onto OPCs. Previous research had found disparate changes when increasing vs. decreasing neuronal activity on both OPC proliferation and myelination(Mitew et al., 2018a) so we sought to examine whether altering neuronal activity (either increasing it or decreasing it, using chemogenetics, chronically, for one week during the period of enhanced developmental synapse formation and myelination, would impact neuron to OPC synapses. We found that increasing neuronal activity alone affected OPC proliferation and OPC activation, and that either increasing or decreasing neuronal activity altered neuron to OPC inhibitory and excitatory synapse formation, as well as the types of neuronal inputs onto OPCs, discussed further below.

## Methods

### Animals

All animal procedures were approved by the Institutional Animal Care and Use Committees at Santa Clara University and the University of California San Francisco. Pdgfra-CreERT2 driver mice were crossed to ROSA-tTA; pTRE-Bi-G-TVA; ROSA-YFP mice and the resulting litters were genotyped in order to generate Pdgfra-CreERT2; ROSA-tTA; pTREBi-G-TVA; ROSA-YFP mice. (Kang et al., 2010) (Hippenmeyer et al., 2005) (Han et al., 2013) (Yasuda & Mayford, 2006)Following tamoxifen-induced gene recombination, Pdgfra-CreERT2; ROSA-tTA; pTREBi-G-TVA; ROSA-YFP mice express the avian envelope protein TVA and the rabies glycoprotein GRab under tetracycline control, as well as a YFP reporter protein, only in OPC cells expressing Pdgfra, allowing for OPC-specific targeting of the viral proteins necessary for infection (TVA) and spread (GRab) along with a label of recombined OPCs. Saline or Clozapine was delivered IP every day for 7 days; Clozapine was delivered at a dose of 0.1mg/kg. Tamoxifen was delivered at a dose of 0.5mg/40g via IP injection was spaced over 7 days with 24 hours between the last dose and either viral injection or euthanasia. Edu was delivered IP every day for 7 days a dose of 10mg/kg.

### Viruses

Two avian envelope protein pseudotyped (EnvA) G deleted mutant rabies viruses that expressed either a red fluorescent reporter ((EnvA)SADΔmCherry) or a green fluorescent reporter ((EnvA)SADΔGFP) were amplified and pseudotyped from stock viruses from the Callaway lab (Salk Institute) following their established protocol. (Wall et al., 2010) Each virus was titered on mammalian 3T3 cells to confirm that the virus expressed the EnvA protein, and to determine the viral titer: 1 × 10^9^ IU ml^-1^ was reached for each round of production.

An AAV virus expressing pAAV-hSyn-hM4D(Gi)-mCherry (AAV8), which would lead to decreased activity in projection neurons of the cortex, was sourced from Addgene (50475), as was an AAV virus expressing pAAV-hSyn-hM3D(Gq)-mCherry (AAV8), which would lead to increased activity in projection neurons of the cortex (Addgene, 50474).

### Viral Circuit Tracing by Brain Region

Pdgfra-CreERT2; ROSA-tTA; pTREBi-G-TVA; ROSA-YFP (N=4-6/group, counterbalanced for sex) were dosed with tamoxifen (DOSE: 2 mg/25 g body weight) via intraperitoneal (IP) injection for 5 days starting from postnatal day 25. 24 hours after the last tamoxifen injection, mice were stereotaxically injected with 300nl of EnvA pseudotyped G deleted rabies virus expressing mCherry (EnvA ΔGRabV-mCherry, 300nl). The stereotaxic injection occurred in CA1 of the hippocampus (X 1.1, Y -1.7, Z 1.4), the dentate gyrus (X 2.1, Y -2.7, Z 2.9), striatum (X 2.5, Y 0, Z 2.6), or cortex (X 0.6, Y -1.5, z 1.0). Following injection, mice recovered and were placed in group housing for a period of 7 days to allow for viral expression, after which they were sacrificed and the brain was collected and processed for histology. Sections were stained with Olig2, NeuN, Parvalbumin, and Somatostatin, and the number of RabV+ cells co-labeled with these stains was quantified as was the total number of RabV+ cells.

### Neuronal Activity Modulation

ROSA-tTA; pTREBi-G-TVA; ROSA-YFP mice (N=4/group, counterbalanced for sex) were stereotaxically injected in the hippocampus with an AAV virus expressing hSyn-hM4D(Gi)-mCherry (400nl, X 1.1, Y -1.7, Z 1.4), or one expressing hSyn-hM3D(Gq)-mCherry (400nl, X 1.1, Y -1.7, Z 1.4) between P8-P12, and returned to group-housing for two weeks to allow for viral expression. Mice were injected with IP injections of EdU (0.1 mg/25g) and IP injections of either saline or clozapine (0.1 mg/kg) for 7 days starting from P25. 24 hours post-injection, mice were sacrificed and the brain was collected and processed for histology. Sections were stained with ctip2 and cfos to confirm viral cell infection specificity. Additional sections were stained with Olig2 and Cfos, and the number of CFos+/Olig2+ cells were quantified for each condition.

### Viral Circuit Tracing Following Neuronal Activity Modulation

Pdgfra-CreERT2; ROSA-tTA; pTREBi-G-TVA; ROSA-YFP (N=4-6/group, counterbalanced for sex, were stereotaxically injected in the hippocampus with an AAV virus expressing hSyn-hM4D(Gi)-mCherry (400nl, X 1.1, Y -1.7, Z 1.4), or one expressing hSyn-hM3D(Gq)-mCherry (400nl, X 1.1, Y -1.7, Z 1.4) between P8-P12, and returned to group-housing for two weeks to allow for viral expression. After two weeks, mice were administered tamoxifen (DOSE: 1 mg/25 g body weight) via IP injection for 5 days as well as an IP injection of clozapine (0.1mg/kg) or saline for 7 days. 24 hours after the last clozapine or saline injection, mice were injected with EnvA ΔGRabV-GFP (300nl, X 1.1, Y -1.7, Z 1.4), recovered, and returned to housing. 7 days after rabies injection, mice were sacrificed and the brain was collected and processed for histology. Sections were stained with Ctip2, Olig2, or Parvalbumin/Somatostatin, and the number of RabV+ cells co-labeled with these stains was quantified as was the total number of RabV+ cells.

### In Vivo Electrophysiology

A microelectrode array (4 × 4 35 µm electrodes with 150 µm row and electrode spacing (Innovative Neurophysiology, Durham, NC) was unilaterally (right hemisphere) targeted to the dHPC (array center: X 1.1, Y -1.7, Z 1.4) and affixed to the skull with dental cement. One week after surgery, mice were habituated to the recording tether for 1 hr/day for 3 days in their home cage prior to recordings.

In vivo electrophysiological recordings took place during 120 min sessions in the home cage. After 10 min of drug-free baseline recording, mice were injected with vehicle or clozapine (0.1mg/kg, IP) and returned to the home cage for the remainder of the recording session. Electrophysiological recordings were acquired using SpikeGadgets main control unit and Trodes software (SpikeGadgets, San Francisco, CA), using 16-channel digitizing head-stages sampled at 30 kHz. Spike sorting was conducted manually using Offline Sorter (Plexon Inc., Dallas, TX) and analyzed using NeuroExplorer (Nex Technologies, Colorado Springs, CO) as previously described (Halladay and Blair, 2015; Halladay et al., 2020).

On completion of testing, mice were anesthetized with 2% isoflurane and a current stimulator (S48 Square Pulse Stimulator, Grass Technologies, West Warwick, RI) delivered 2 seconds of 40 µA DC current through each electrode to make a small marking lesion. 24 hr later, mice were overdosed via 150 mg/kg IP Euthasol (Henry Schein, Melville, NY) and perfused intracardially with PBS followed by 4% PFA. Brains were left in 4% PFA overnight, then transferred to a 30% sucrose PBS solution for cryoprotection. Coronal sections (50 µm thick) were cut on a cryostat (Leica Biosystems Inc, Buffalo Grove, IL) and mounted onto slides. Tissue was stained with DAPI (Sigma-Aldrich) and imaged using a Keyence BZ-X800 fluorescence microscope (Keyence Corporation of America, Itasca, IL). Only single units with placements confirmed to be in the dHPC were included in the analysis.

To determine whether units altered their firing rate following vehicle or clozapine administration, data during 60-sec bins following the injection were z-score normalized to a 10-min baseline period just prior to injection. For individual units, time bins with a z-score of > |1.96| were considered significantly different than baseline (p<.05) (Figure ** [color maps]). Results from one-way ANOVA confirmed that unit firing rate was significantly dependent upon virus type (F_2,357_= 330.23, p<.0001). Planned post hoc comparisons of averaged viral group unit responses (independent samples t-tests) revealed that compared to the control group, units recorded in mice injected with hM3Dq and hM4Di viral constructs displayed significantly greater or lesser, respectively, average normalized firing rates during the recording session, beginning 15 and 9 minutes after clozapine administration (p<.01). [each significant bin denoted by orange/blue lines at top of right panel graph]

### Tissue Processing and Sectioning

Mice were sacrificed via transcardial perfusion under terminal anesthesia with sodium pentobarbital (250 mg/kg). Anesthetized mice were pinned and visualization of the heart was achieved following removal of the rib cage, after which a 21-gauge needle was inserted into the left ventricle, the vena cava snipped, and 10 ml of ice-cold PBS run through the line, followed by 10 ml of 4% paraformaldehyde. After perfusion, the brain was harvested and placed at 4ºC in 4% PFA for 2 hours, after which it was moved to 15% sucrose overnight, and then cryoprotected in 30% sucrose overnight. The brain was then embedded in O.C.T. compound (Sakura Finetek; Torrance, CA, USA) on dry ice and stored at -80 ºC until sectioned into 25 µm slices on a CM1860 Leica cryostat (Leica Microsystems).

### Immunohistochemistry

After sections were washed thrice with PBST for 10 minutes at a time, they were incubated in blocking buffer (10% lamb serum, 0.33% Triton, 0.005% sodium azide, and 1:50000 DAPI in PBS) for 1 hour, after which they were incubated in primary antibody for 4 hours at room temperature. Following three washes with PBST, sections were then incubated in secondary antibody for 1 hour, followed by three washes in PBST, and mounted using Fluoromount-G mounting medium (Thermo Fisher Scientific; Waltham, MA, USA). Stained sections were imaged using a Zeiss Axioscanner, and representative images for each condition were collected using a Zeiss 710 confocal microscope. The following primary antibodies were used: anti-Olig2 Rb (Abcam ab109186); anti-Olig2 Ms (Millipore MAB N50), anti-Parvalbumin Ms (Millipore MAB1572), anti-Somatostatin Rb (Invitrogen PA585759), anti-Ctip2 Rat (Abcam AB18465), anti-CFos Rb (Synaptic Systems 226 008), anti-Gephyrin Ms (Synaptic Systems 147021), anti-SAP97 (Thermofisher PA1-741), and anti-NeuN Ms (Thermofisher PA5-78499). All secondary antibodies were raised in goat and were Alexa Fluor secondary antibodies from Jackson Immunoresearch, conjugated to 488, 546, or 633 fluorophores as needed.

### EdU Staining

After sections were washed thrice with PBST for 10 minutes at a time, they were incubated in blocking buffer (10% lamb serum, 0.33% Triton, 0.005% sodium azide, and 1:50000 DAPI in PBS) for 1 hour, after which they were stained with EdU using the Click-iT® EdU Imaging Kit with Alexa Fluor® 647 Azide (ThermoFisher, C10337, Waltham, MA, USA) according to manufacturer instructions. Following EdU staining, slides were immunostained with Olig2. Slides were washed in PBST three times and mounted using Fluoromount-G mounting medium. Representative images for each condition were collected using a Zeiss 710 confocal microscope. The number of EdU+/Olig2+ cells in the hippocampus were quantified for each condition.

### Data Collection and Statistical Analyses

Sections were analyzed in order to manually quantify the number of co-labeled cells by category. Zeiss Zen lite was used to visualize the image files from the Zeiss Axioscanner for quantification.

Quantification of Olig2 cells followed the processing of imaging by subtracting background, adjusting the threshold, and processing the result on the Binary menu with watershed and making mask. Then we finally quantified particles on the sizes of 10um and roundness of 0.1 um.

In order to quantify gephyrin and SAP97 we measured the corrected total cell fluorescence (CTCF) using the formula: CTCF = Integrated Density – (Area of selected cell X Mean fluorescence of background readings). First, we sampled the cells of interest randomly on Olig2 labeled channel along CA1 and DG regions, using drawing/selection tool. Adjacent to the cells other areas were selected to be used as background measurement. Using the “measure” function we collected area, integrated density, and mean grey values on the Gephyrin and SAP97 channels and applied the CTCF formula.

Statistical analyses were performed using unpaired t tests for the Gq and Gi conditions separately using GraphPad Prism version 8.0.0.

## RESULTS

Based on previous work (Spitzer et al., 2019), we hypothesized that there would be regional differences in neuron to OPC connections, with brain regions having unique patterns of connectivity between OPCs, both in terms of the cytoarchitecture of the connections, the number of connections, and the types of presynaptic inputs onto OPCs. Given the neuron to OPC connections are transient and are lost when OPCs differentiate into oligodendrocytes (de Biase et al., 2010), we used an inducible Pdgfra-Cre mouse to label OPCs during postnatal development, at the peak of OPC differentiation and myelination, when we hypothesized there would be large numbers of neuron to OPC connections. We targeted different regions of the forebrain with unique cytoarchitecture, circuitry, and myelin patterns, to examine the diversity of these connections.

Representative images of labeled presynaptic neurons from the somatosensory cortex, dorsal striatum, CA1 of the hippocampus, and dentate gyrus illustrate several interesting and diverse patterns of neuron to OPC connections. Long distance cortical inputs to OPCs are primarily pyramidal neurons with few contralateral inputs, whereas CA1 OPCs receive many contralateral inputs from hippocampal pyramidal neurons and cortical connections: DG targeting reveals CA1 and CA3 neuronal inputs. Striatal targeting is mainly limited to local striatal projection neuron inputs with rare contralateral connections or inputs outside of the striatum (Figure 1, B-M). Further, the patterns of these connections were consistent with their local circuitry: cortical connections were often columnar, with inputs stretching across many layers of the cortex; striatal connections were largely nuclear, with scattered inputs within the caudate and putamen; inputs to CA1 localized OPCs were from many neurons throughout the stratum pyramidal of CA1 and the stratum lucidum of CA3; DG inputs to OPCs were made up of both local granule cells and mossy cells as well as occasional neurons in the entorhinal cortex (Figure 1, B-M; data not shown). We also examined whether both excitatory and inhibitory neurons made similar numbers of inputs onto OPCs in these different regions. (Figure 1N). Interestingly, there were regional differences in the number of neuronal inputs onto OPCs, with the OPCs in the hippocampus receiving relatively more neuronal inputs, compared to the striatum and cortex (Figure 1O). Interestingly we found that the CA1 of the hippocampus had the fewest PV+ or SST+ inhibitory neuron (IN) synapses onto OPCs, followed by the DG. The cortex and striatum both had many more INs connected to OPCs than in the hippocampus, and there were more PV+ INs than SST INs in the cortex, whereas in the striatum there were more SST+ INs than PV+ INs. Finally, we found regional differences in the number of neuron to OPC synapses, which were much higher in the CA1 of the hippocampus compared to the other three regions. The patterns of connectivity were also regionally unique: cortical OPCs received synapses across many layers of the cortex, with no layer restrictions to inputs. However, cortical inputs were often columnar in structure, spanning the cortex, but not invading adjacent columns. In the hippocampus, CA1 injections largely yielded CA1 or CA3 pyramidal neuron inputs; these were sometimes only ipsilateral but often contralateral as well. DG injections yielded mostly local granule cell labeling; this was infrequently bilateral. Striatum labeled neurons were a mix of NeuN+ neurons and PV+ or SST+ INs, but also followed the local nuclear structure of the striatum and were rarely bilateral. The diversity of connectivity between neurons and OPCs in the forebrain, as well as the high level of neuronal inputs onto each OPC in the hippocampus led us to focus our future experiments on the CA1 region of the hippocampus.

**Fig 1.**
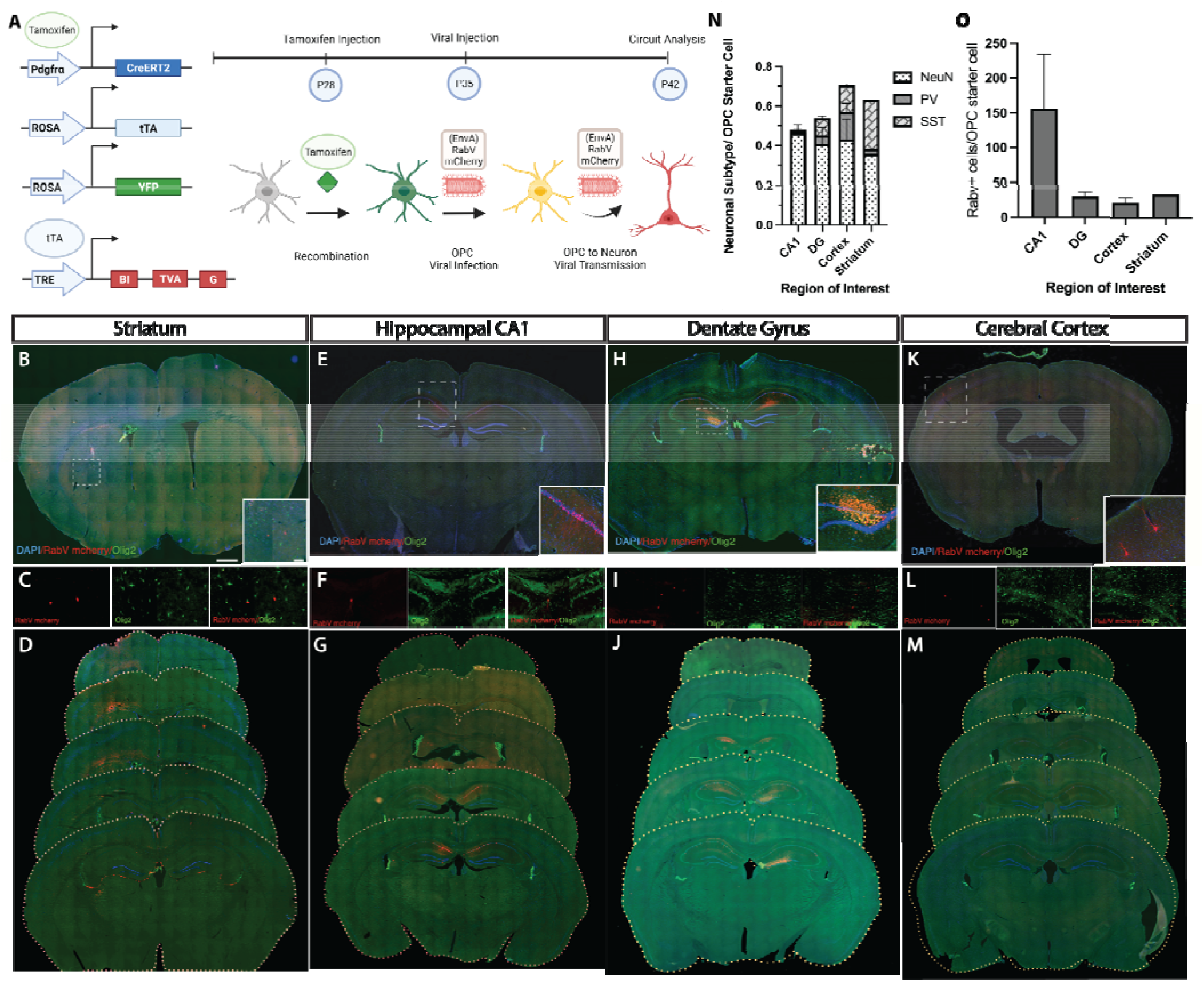
Neuron to OPC Circuits Have Diverse Patterns of Connectivity in the Forebrain. A. We combined attenuated rabies virus with a Cre-loxP based system using the Pdgfra-^CreERT2^ driver mouse crossed to ROSA^tTA;^ pTRE-Bi-^G-TVA^; tau-^mGFP^ mice, inducing recombination in OPCs using tamoxifen. These mice now express the avian envelope protein TVA and the rabies glycoprotein GRab as well as a GFP reporter protein. This allows us to target only OPCs with the viral proteins necessary for infection (TVA) and spread (GRab) along with a label of recombined OPCs. We target OPCs with a viral injection of avian envelope protein pseudotyped deletion mutant rabies virus (EnvA)RabV that expresses a red fluorescent reporter (mCherry), prepared as in (Wickersham, et al, Nature methods). (B-D) Striatum, (E-G) CA1, (H-J) Dentate Gyrus, and (K-M) Somatosensory Cortex targeting of neuron to OPC circuits using monosynaptic viral circuit tracing. Pdgfra-CreERT2; YFP; tTA; G-TVA mice injected at P60 with (EnvA)RabV mcherry after 5 days of tamoxifen treatment. (C,F,I,L) Neurons (labeled in red) are presynaptically connected to each OPC population (labeled with both red and green). (N) Quantification of: neuronal (Neun+) inputs, PV+ interneuron inputs, and SST+ interneuron inputs connected to OPCs in CA1, dentate, striatum, and somatosensory cortex. (O) Quantification of number of RabV+ inputs per OPC starter cell in each of the brain regions in B-M. Scale bar: 50um, 500um.

We used chemogenetics to manipulate neuronal activity in vivo with the goal of targeting CA1 neurons in the hippocampus. We confirmed that injection of pAAV-hSyn-hM3D(Gq)-mCherry or pAAV-hSyn-hM4D(Gi)-mCherry into the CA1 of the hippocampus and subsequent injection of saline did not induce activation of Ctip2+ pyramidal neurons of the hippocampus (Figure 2 A-E). Further, injection of pAAV-hSyn-hM4D(Gi)-mCherry into the CA1 of the hippocampus and subsequent injection of clozapine IP for 7 days resulted in no activation of Ctip2+ pyramidal neurons of the hippocampus (Figure 2 F,G). However, injection of pAAV-hSyn-hM3D(Gq)-mCherry into the CA1 of the hippocampus and subsequent injection of clozapine IP for 7 days resulted in activation of Ctip2+ pyramidal neurons of the hippocampus, resulting in robust c-fos expression in these neurons (Figure 2 H,I). To confirm this effect in real time, and to ensure that the pAAV-hSyn-hM4D(Gi)-mCherry injection with clozapine resulted in neuronal inhibition, as well as to confirm that there were no baseline effects of the DREADDs or clozapine alone, we used conducted in vivo array recordings to measure the changes in firing rate in individual unit recordings in the CA1 of the hippocampus. (Figure 2J). We found that saline in pAAV-hSyn-hM4D(Gi)-mCherry or saline in pAAV-hSyn-hM3D(Gq)-mCherry injected mice had no effect on neuronal spiking (top left and middle histograms), nor did clozapine treatment alone (bottom histograms), but a 7 day clozapine treatment in pAAV-hSyn-hM4D(Gi)-mCherry mice resulted in inhibition of neuronal activity in the hippocampus, and a 7 day clozapine treatment in pAAV-hSyn-hM3D(Gq)-mCherry injected mice resulted in robust activation of neuronal activity in the hippocampus (top right and middle histograms and Figure 2K).

**Fig 2.**
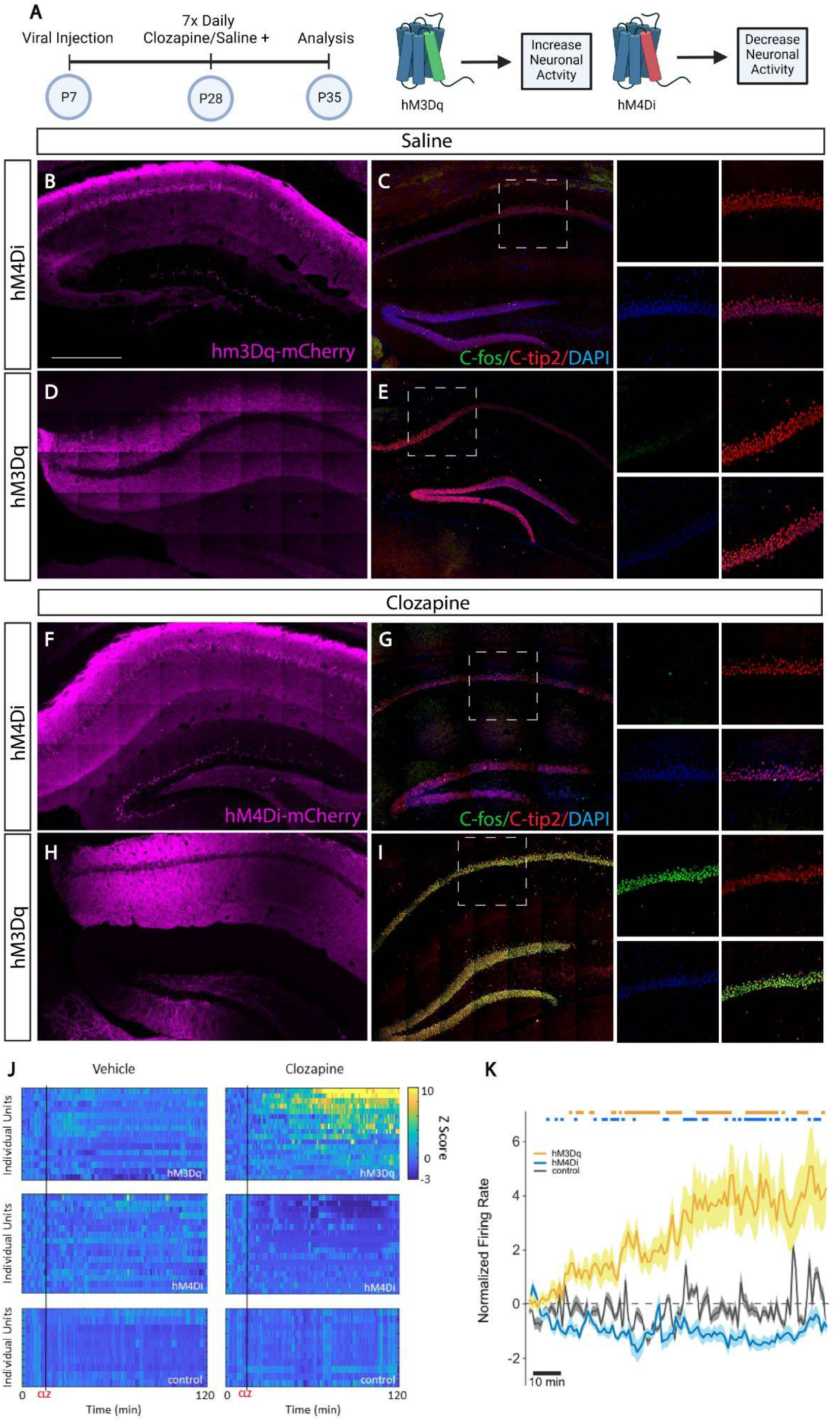
DREADD (Designer Receptors Exclusively Activated by Designer Drugs) Activator Gq Results in Activation and DREADD Activator Gi Results in Inhibition of the Hippocampal Neuronal Circuit. (A) pAAV-hSyn-HA-hM3D(Gq)mCherry or pAAV-hSyn-hM4D(Gi)-mCherry virus were injected into the CA1 of the hippocampus at P7. 3 weeks later, animals were treated with 7 daily IP doses of clozapine or saline. Animals were euthanized 24 hours after the last clozapine dose at P35. (B, F) Representative expression of hM4D(Gi)mCherry in the hippocampus. (D,H) Representative expression of hM3D(Gq)mCherry in the hippocampus. (C) Saline treatment in hM4D(Gi)mCherry expressing mice or (E) Saline treatment in hM3D(Gq)mCherry expressing mice results in no c-fos labeling of ctip2+ neurons. (G) 7-day Clozapine treatment in hM4D(Gi)mCherry expressing mice resulted in very low cfos labeling of ctip2+ neurons. (H) G) 7-day Clozapine treatment in hM3D(Gq) mCherry expressing mice resulted in very high cfos labeling of ctip2+ neurons. (J) In vivo array recording of cells in CA1 of the hippocampus of mice injected with IP Clozapine or Saline. Top histograms: Representative cells recorded from the hippocampus of littermates infected with hM3Gq) and injected with saline (left) or clozapine (right). Middle histograms: representative cells recorded from the hippocampus of littermates infected with hM4D(Gi) and injected with saline (left) or clozapine (right). Bottom histograms: representative cells recorded from the hippocampus of littermates treated with clozapine but no hM4D(Gi) injection (bottom left) or treated with clozapine but no hM3D(Gq) injection (bottom right). K. Average suppression rates from cells in the hippocampus of mice on the 7th day of Clozapine treated hM3D(Gq), Clozapine treated hM4D(Gi), or Saline treated control mice. Scale bar: 500um.

Once we established that we had a robust approach to increase or decrease neuronal activity, we examined the effect of chronically increased neuronal activity or chronically decreased neuronal activity on OPC development, measuring the number of dividing cells with time-matched Edu and either Clozapine or saline treatment. We hypothesized that decreasing neuronal activity would result in fewer OPCs proliferating, as neuronal activity has been shown to be an important cue for OPC proliferation and differentiation (Bergles et al., 2010)(Gallo et al., 1996), (Yuan et al., 1998), (Gudz et al., 2006). We found this not to be the case: there were more dividing OPCs in the hM4D(Gi) mice treated with clozapine, compared to saline treatment alone (Figure 3B-C). Additionally, we hypothesized that, consistent with previous work (Mitew et al., 2018b), increasing neuronal activity would result in increased proliferation of OPCs. We found this to be the case: increasing neuronal activity resulted in an increase in the number of dividing OPCs, compared to saline treatment alone, although this was a trend and not a significant effect in our data, due to variability in our Gq clz treatment group (Figure 3D-E). The increase in proliferation was consistent with previous work, although others have not shown increased proliferation after neuronal activity. This indicates that there is a larger pool of OPCs available after increasing or decreasing neuronal activity. We were curious as to whether this might be a mechanism to induce plasticity in the OPCs themselves, and so we stained them with c-fos, an immediate early gene that is used to measure plasticity in neural cells (Kovacs et al., 2008). We found that either increasing or decreasing neuronal activity led to increases in the expression of the plasticity gene c-fos in OPCs, consistent with the idea that altering neuronal activity has functional consequences in OPCs (Figure 4). This effect was specific to the clozapine treated animals, and was not found in saline treated controls.

**Figure 3.**
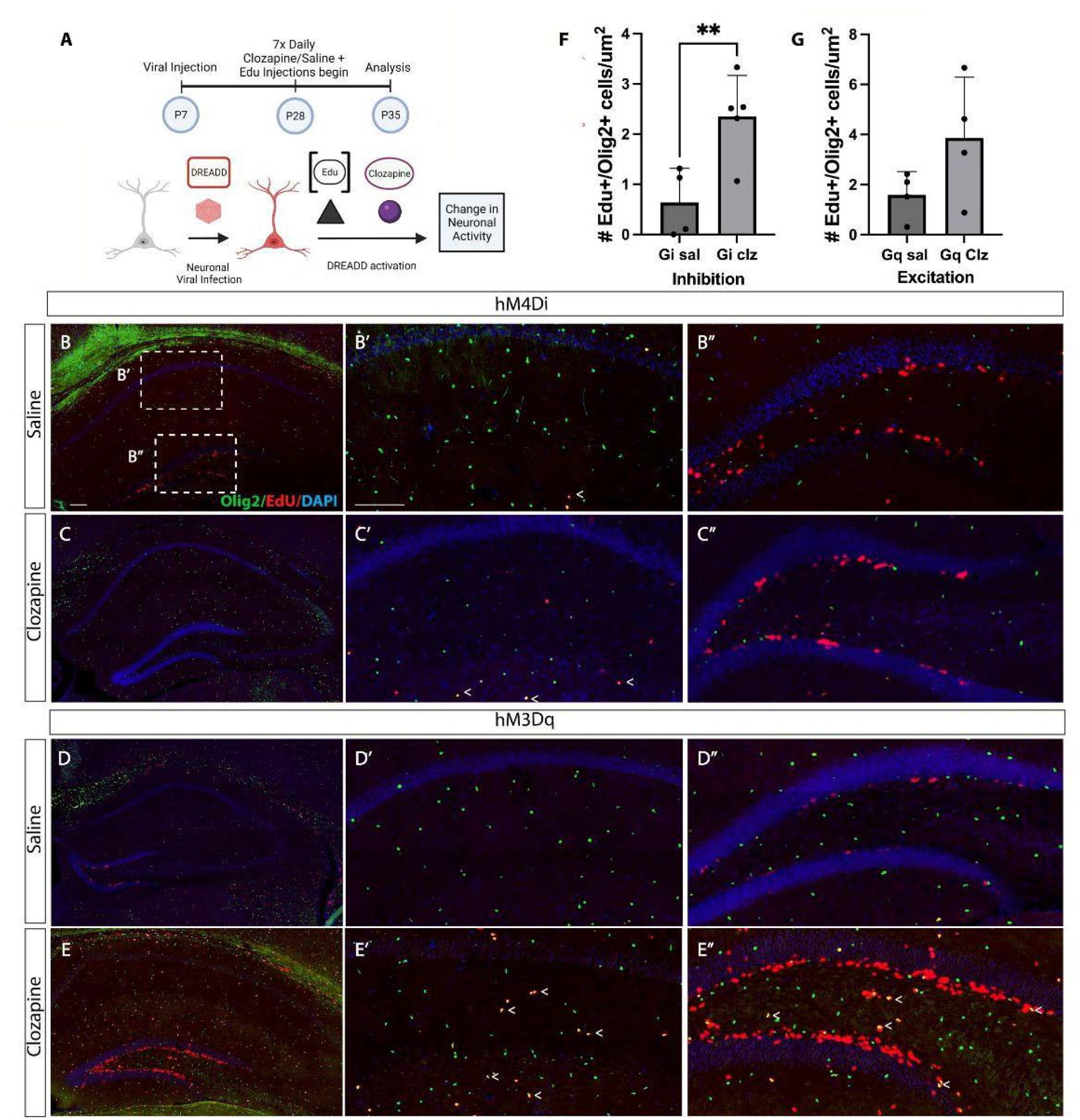
Altering Neuronal Activity Leads to Increased Cell Proliferation in the Hippocampus. (A) pAAV-hSyn-HA-hM3D(Gq)mCherry or pAAV-hSyn-hM4D(Gi)-mCherry AAV was injected into the CA1 of the hippocampus at P7. 3 weeks later, animals were treated with 7 daily IP doses of Edu plus Clozapine or Saline. Animals were euthanized 24 hours after the last dose at P35. Confocal images of Edu+ olig2+ OPCs after: (B) Saline treatment in hM4Di injected animals, (C) Clozapine treatment in hM4Di injected animals, (D) Saline treatment in hM3Dq injected animals, (E) Clozapine treatment in hM3Dq injected animals, (F) Ratio of dividing OPCs/total OPCs in hM4Di injected animals (G) Number of dividing cells/um^2^ in hM4Di injected animals (H) Ratio of dividing OPCs/total OPCs in hM3Dq injected animals (I) Number of dividing cells/um^2^ in hM3Dq injected animals. Scale bar: 100um.

**Fig 4.**
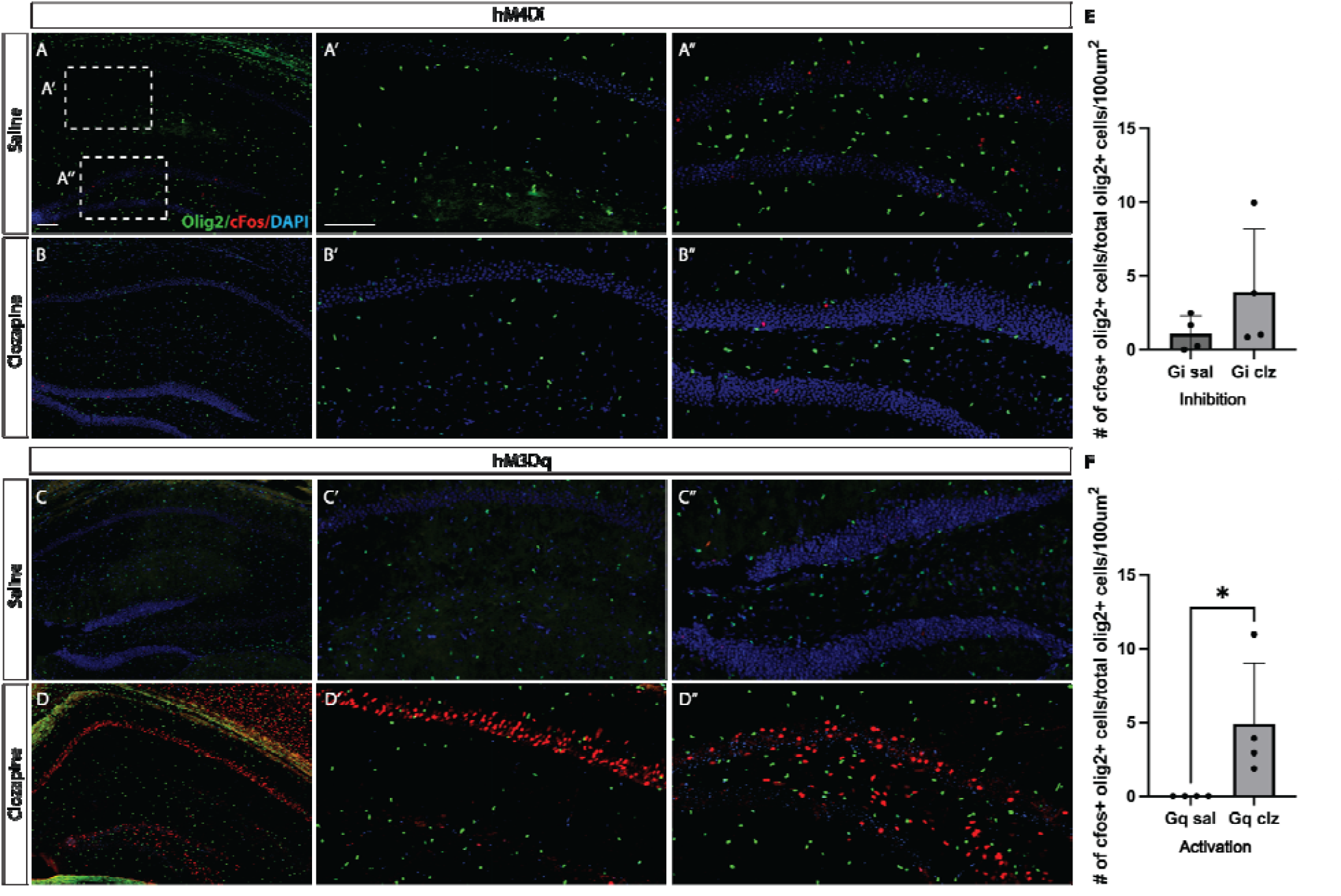
Increasing Neuronal Activity Increases Direct Activation of OPCs. pAAV-hSyn-HA-hM3D(Gq)mCherry or pAAV-hSyn-hM4D(Gi)-mCherry AAV was injected into the CA1 of the hippocampus at P7. 3 weeks later, animals were treated with 7 daily IP doses of Clozapine or Saline. Animals were euthanized 24 hours after the last dose at P35. Confocal images of cfos+/olig2+ OPCs after: (A) Saline treatment in hM4Di injected animals, (B) Clozapine treatment in hM4Di injected animals, (C) Saline treatment in hM3Dq injected animals, (D) Clozapine treatment in hM3Dq injected animals, (E) Ratio of activated OPCs/total OPCs in hM4Di injected animals (F) Ratio of activated OPCs/total OPCs in hM3Dq injected animals. Scale bar: 100um.

To determine the functional consequences of increased activity on OPCs, we next examined whether synapse formation onto OPCs was also affected. We used the same chemogenetic approach to increase or decrease neuronal activity, and examined the changes in the expression of the postsynaptic density protein Gephyrin as a readout of the formation of inhibitory neuronal synapses onto OPCs. We hypothesized that decreasing neuronal activity would result in increased Gephyrin positive puncta onto OPCs, as silencing all neuronal synapses would result in compensation by increasing the number of inhibitory synapses. As shown in Figure 5 I-J, we found that the number of Gephyrin+ puncta on OPCs increased in hM4D(Gi) mice treated with clozapine. This was true in CA1 but not in the DG of the hippocampus, indicating this plasticity at neuron to OPC synapses was local. Furthermore, we hypothesized that increasing neuronal activity would lead to increased Gephyrin expression, as a compensatory mechanism for OPCs to regulate the increased neuronal activity by adding more inhibitory synapses. As shown in Figure 5 K-L, we found that the number of Gephyrin+ puncta on OPCs increased in hM3D(Gq) mice treated with clozapine. This was true in both CA1 and in the DG of the hippocampus, indicating this plasticity at neuron to OPC synapses was present throughout the hippocampal network.

**Figure 5.**
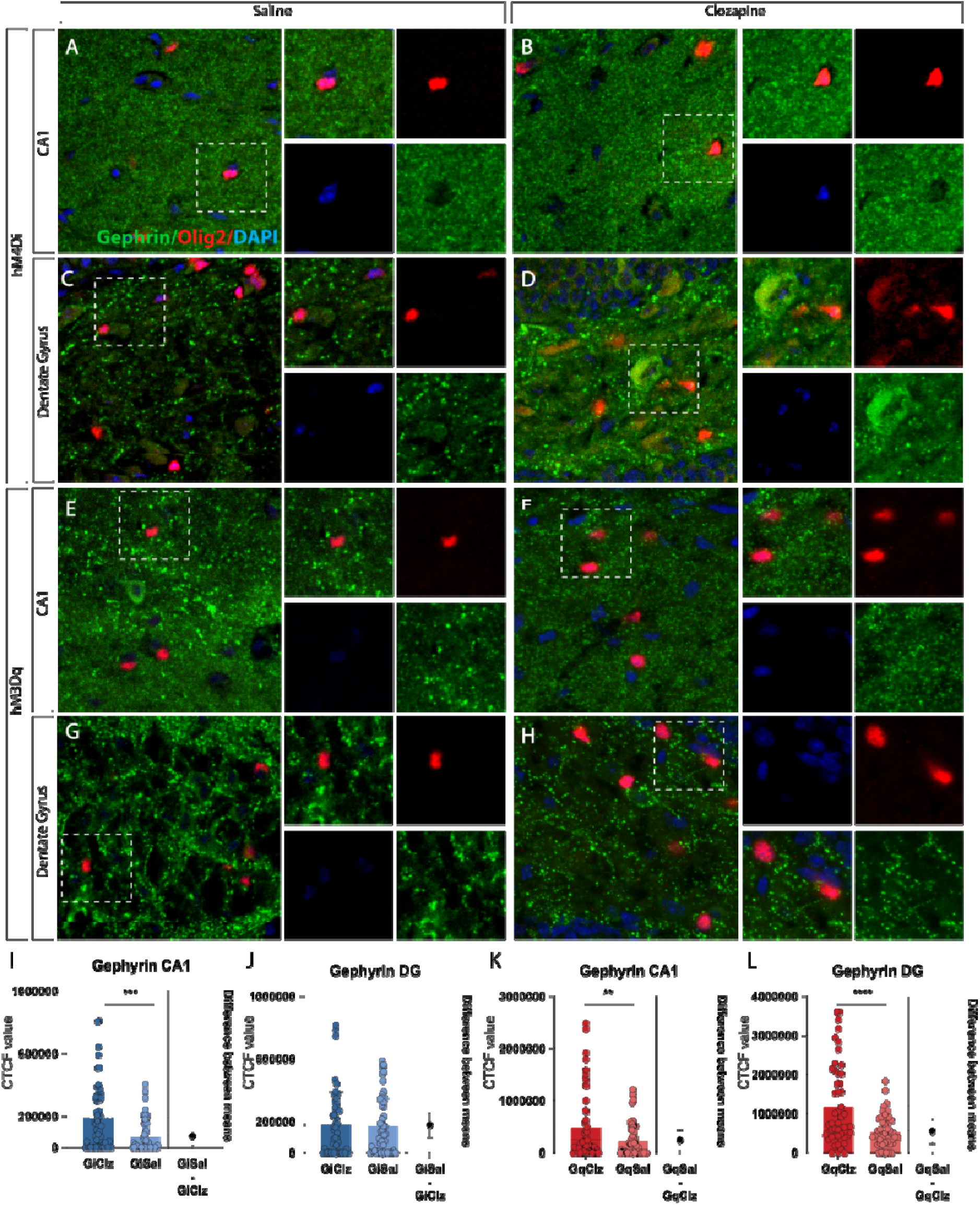
Manipulation of Neuronal Presynaptic Activity onto OPCs Changes Inhibitory Synapses on OPCs. pAAV-hSyn-HA-hM3D(Gq)mCherry or pAAV-hSyn-hM4D(Gi)-mCherry AAV was injected into the CA1 of the hippocampus at P7. 3 weeks later, animals were treated with 7 daily IP doses of Clozapine or Saline. Animals were euthanized 24 hours after the last dose at P35. Confocal images of Gephyrin+ Olig2+ OPCs after: Saline treatment in hM4Di injected animals in (A) CA1 and (C) DG, Clozapine treatment in hM4Di injected animals in (B) CA1 and (D), Saline treatment in hM3Dq injected animals in (E) CA1 and (G) DG, or Clozapine treatment in hM3Dq injected animals in (F) CA1 and (H) DG. (I) CTCF value of gephyrin puncta in OPCs in hM4Di injected animals in CA1 (J) CTCF value of gephyrin puncta in OPCs in hM4Di injected animals in DG (K) CTCF value of gephyrin puncta in OPCs in hM3Dq injected animals in CA1 (L) CTCF value of gephyrin puncta in OPCs in hM3Dq injected animals in DG. Scale bar: 100um.

We used the same chemogenetic approach to increase or decrease neuronal activity, and examined the changes in the expression of the postsynaptic density protein Gephyrin as a readout of the formation of inhibitory neuronal synapses onto OPCs (Figure 5 A-H). We hypothesized that decreasing neuronal activity would result in increased gephyrin positive puncta onto OPCs, as silencing all neuronal synapses would result in compensation by increasing the number of inhibitory synapses. As shown in Figure 5 I-J, we found that the number of Gephyrin+ puncta on OPCs increased in hM4D(Gi) mice treated with clozapine. This was true in CA1 but not in the DG of the hippocampus, indicating that plasticity at neuron to OPC synapses was local. Furthermore, we hypothesized that increasing neuronal activity would lead to increased Gephyrin expression, as a compensatory mechanism for OPCs to regulate the increased neuronal activity by adding more inhibitory synapses. As shown in Figure 5 K-L, we found that the number of Gephyrin+ puncta on OPCs increased in hM3D(Gq) mice treated with clozapine. This was true in both CA1 and in the DG of the hippocampus, indicating that plasticity at neuron to OPC synapses was present throughout the hippocampal network. We next examined the consequences of manipulating neuronal activity on excitatory synapses.

We again increased or decreased neuronal activity, and examined the changes in the expression of the postsynaptic density protein SAP97 as a readout of the formation of excitatory neuronal synapses onto OPCs (Figure 6A-H). We hypothesized that decreasing neuronal activity would result in increased SAP97+ puncta onto OPCs, as silencing all neuronal synapses would result in compensation by increasing the number of excitatory synapses. Surprisingly, as shown in Figure 6 I-J, we found that the number of SAP97+ puncta on OPCs decreased in hM4D(Gi) mice treated with clozapine compared to controls, indicating a loss of excitatory synapses onto OPCs after neuronal silencing. This was true in CA1 but not in the DG of the hippocampus, indicating that plasticity at neuron to OPC synapses was local. Furthermore, we hypothesized that increasing neuronal activity would lead to decreased SAP97 expression, as a compensatory mechanism for OPCs to regulate the increased neuronal activity by removing excitatory synapses. As shown in Figure 6 K-L, we found that the number of SAP97+ puncta on OPCs decreased in hM3D(Gq) mice treated with clozapine compared to controls. This was true in both CA1 and in the DG of the hippocampus, indicating that plasticity at neuron to OPC synapses was present throughout the hippocampal network.

**Figure 6.**
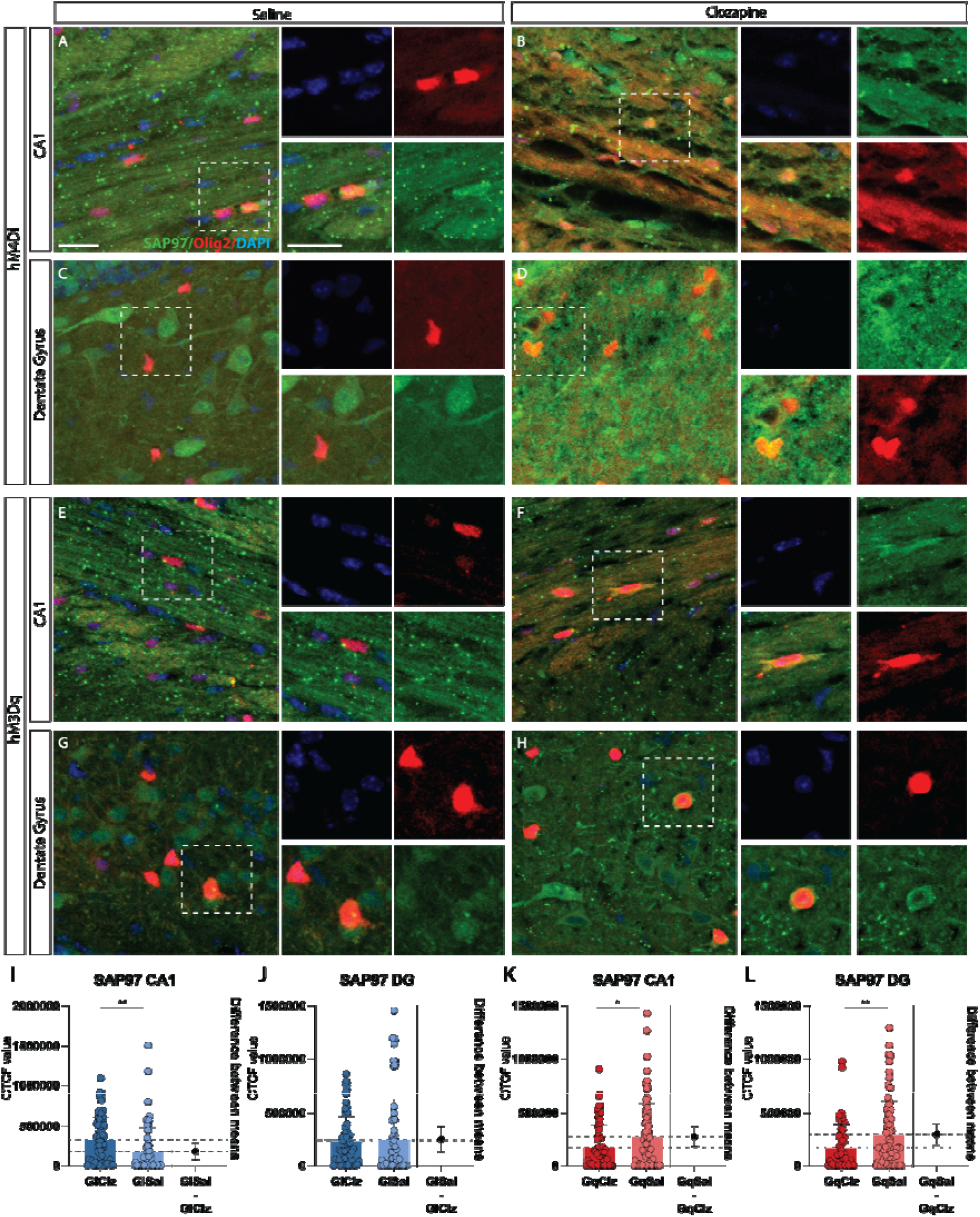
Manipulation of Neuronal Presynaptic Activity onto OPCs Changes Excitatory Synapses on OPCs. pAAV-hSyn-HA-hM3D(Gq)mCherry or pAAV-hSyn-hM4D(Gi)-mCherry AAV was injected into the CA1 of the hippocampus at P7. 3 weeks later, animals were treated with 7 daily IP doses of Clozapine or Saline. Animals were euthanized 24 hours after the last dose at P35. Confocal images of SAP97+ Olig2+ OPCs after: Saline treatment in hM4Di injected animals in (A) CA1 and (C) DG, Clozapine treatment in hM4Di injected animals in (B) CA1 and (D), Saline treatment in hM3Dq injected animals in (E) CA1 and (G) DG, or Clozapine treatment in hM3Dq injected animals in (F) CA1 and (H) DG. (I) CTCF value of SAP97 puncta in OPCs in hM4Di injected animals in CA1 (J) CTCF value of SAP97 puncta in OPCs in hM4Di injected animals in DG (K) CTCF value of SAP97 puncta in OPCs in hM3Dq injected animals in CA1 (L) CTCF value of SAP97 puncta in OPCs in hM3Dq injected animals in DG. Scale bar: 100um.

After finding these changes in excitatory and inhibitory synapse formation onto OPCs, we wanted to determine whether there were global changes in connectivity between neurons and OPCs after chronic manipulation of neuronal activity. We combined chemogenetic manipulations of neuronal activity with genetically targeted monosynaptic tracing using deletion mutant rabies virus to determine the changes in presynaptic inputs onto OPCs (Figure 7A). We found that increasing or decreasing neuronal activity did not change the total number of neuronal inputs onto OPCs (Figure 7). We examined the number of neuronal inputs by region, by local vs. long-distance connections, and by ipsilateral or contralateral inputs, and found no differences (data not shown).

**Figure 7.**
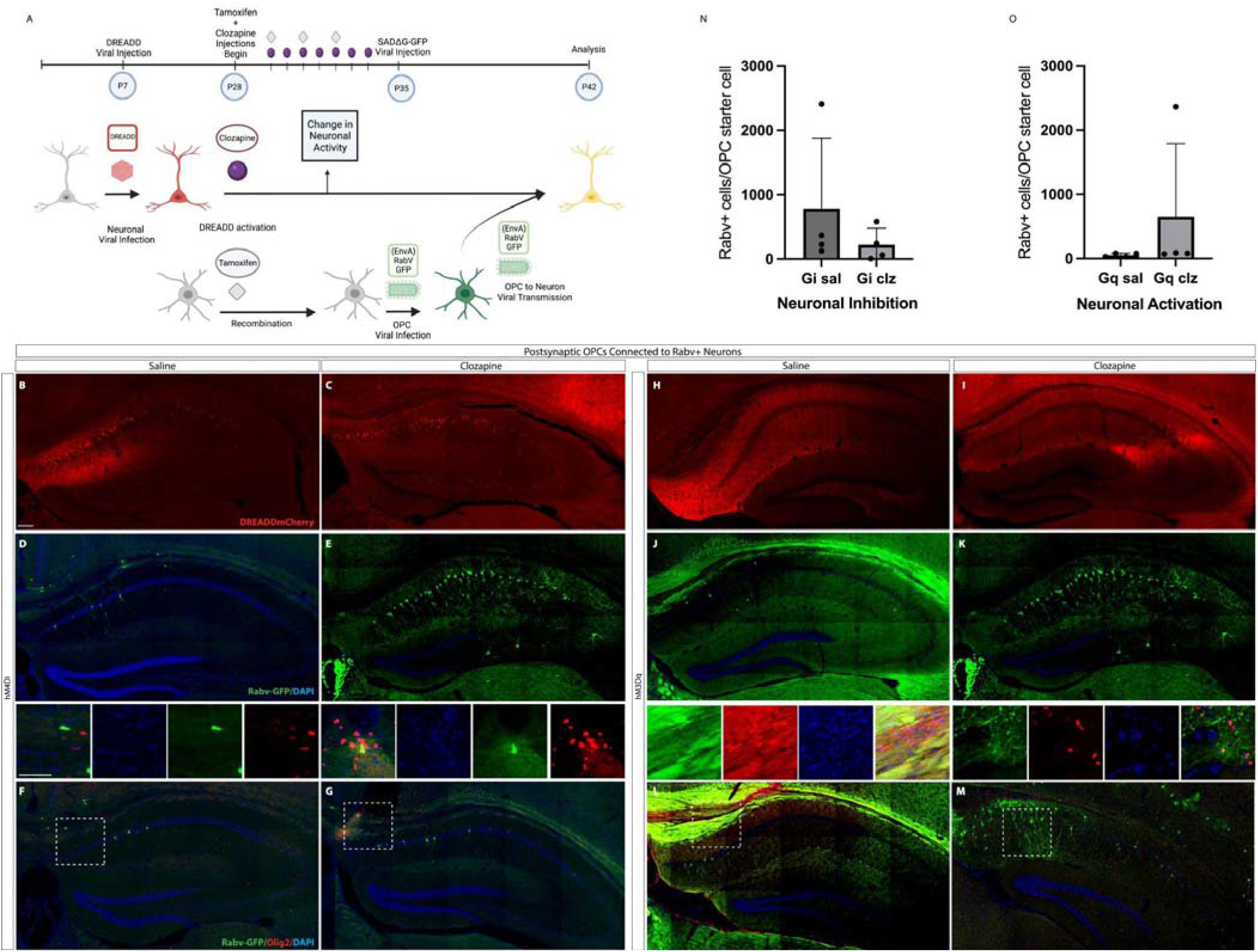
Manipulation of Neuronal Presynaptic Activity onto OPCs Changes Neuron to OPC Connectivity. (A) pAAV-hSyn-HA-hM3D(Gq)mCherry or pAAV-hSyn-hM4D(Gi)-mCherry AAV was injected into the CA1 of the hippocampus at P7. 3 weeks later, animals were treated with 7 daily IP doses of Clozapine or Saline and 4 doses of tamoxifen in that 7-day period. (B) 24 hours a after the last injection, Pdgfra-^CreERT2^;ROSA^tTA;^ pTRE-Bi-^G-TVA^; tau-^mGFP^ mice, inducing recombination in OPCs using tamoxifen. We then targeted OPCs with in CA1 a viral injection of (EnvA)RabV that expresses a green fluorescent reporter (GFP) to measure changes in connections between neurons and OPCs after manipulation of neuronal activity. (B) Red, hM4D(Gi)-mCherry AAV injected in mice treated with saline, (C) Red, hM4D(Gi)-mCherry AAV injected in mice treated with clozapine (D) Green/Blue, (EnvA)RabV-GFP/DAPI in hM4D(Gi) injected mice treated with saline, (E) Green/Blue, (EnvA)RabV-GFP/DAPI in hM4D(Gi) injected mice treated with clozapine (F) Different section, Green/Blue/Red (EnvA)RabV-GFP/DAPI/Olig2 in hM4D(Gi)-injected in mice treated with saline, (G)Different section, Green/Blue/Red (EnvA)RabV-GFP/DAPI/Olig2 in hM4D(Gi)-injected in mice treated with Clozapine. (H) Red, hM3D(Gq)-mCherry AAV injected in mice treated with saline, (I) Red, hM3D(Gq)-mCherry AAV injected in mice treated with clozapine (J) Green/Blue, (EnvA)RabV-GFP/DAPI in hM3D(Gq) injected mice treated with saline, (K) Green/Blue, (EnvA)RabV-GFP/DAPI in hM3D(Gq) injected mice treated with clozapine (L) Different section, Green/Blue/Red (EnvA)RabV-GFP/DAPI/Olig2 in hM3D(Gq)-injected in mice treated with saline, (M)Different section, Green/Blue/Red (EnvA)RabV-GFP/DAPI/Olig2 in hM3D(Gq)-injected in mice treated with Clozapine. (N-P) Scale bar (B) 100um, (D) inset, 50um.

**Figure 8.**
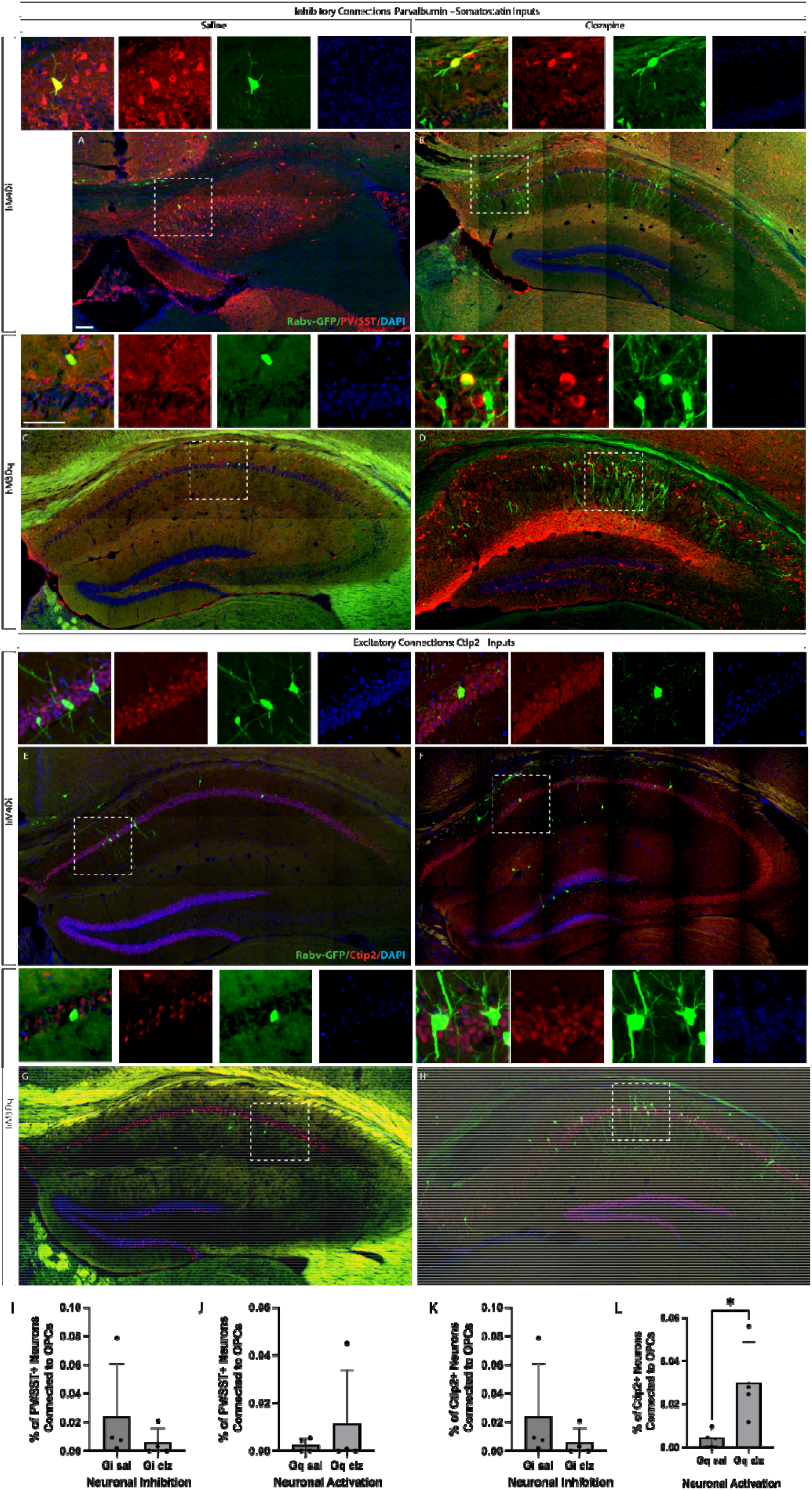
Manipulation of Neuronal Activity onto OPCs Changes Presynaptic Neuronal Inputs Onto OPCs. pAAV-hSyn-HA-hM3D(Gq)mCherry or pAAV-hSyn-hM4D(Gi)-mCherry AAV was injected into the CA1 of the hippocampus, treated with 7 daily IP doses of Clozapine or Saline, 4 doses of tamoxifen in Pdgfra-^CreERT2^;ROSA^tTA;^ pTRE-Bi-^G-TVA^; tau-^mGFP^ mice and then injected with (EnvA)RabV(GFP) to measure changes in connections between neurons and OPCs. (A) Green/Blue/Red (EnvA)RabV-GFP/DAPI/PV/SST in hM4D(Gi)-injected in mice treated with saline, (B) Green/Blue/Red (EnvA)RabV-GFP/DAPI/PV/SST in hM4D(Gi)-injected in mice treated with Clozapine. (C) Green/Blue/Red (EnvA)RabV-GFP/DAPI/PV/SST in hM3D(Gq)-injected in mice treated with saline, (D) Green/Blue/Red (EnvA)RabV-GFP/DAPI/PV/SST in hM3D(Gq)-injected in mice treated with Clozapine. (E) Green/Blue/Red (EnvA)RabV-GFP/DAPI/Ctip2 in hM4D(Gi)-injected in mice treated with saline, (F) Green/Blue/Red (EnvA)RabV-GFP/DAPI/Ctip2 in hM4D(Gi)-injected in mice treated with Clozapine. (G) Green/Blue/Red (EnvA)RabV-GFP/DAPI/Ctip2 in hM3D(Gq)-injected in mice treated with saline, (H) Green/Blue/Red (EnvA)RabV-GFP/DAPI/Ctip2 in hM3D(Gq)-injected in mice treated with Clozapine. Scale bar (A) 100um, (C) inset, 50um.

We hypothesized that the total number of neuronal inputs might be unaffected, but the specific type of input would be altered by neuronal activity. We hypothesized that silencing neuronal inputs would lead to a decrease in the number of excitatory inputs to OPCs, but no change in the number of inhibitory inputs to OPCs. We found this not to be the case: the number of inhibitory inputs to OPCs did not change with a decrease in neuronal activity. We also hypothesized that increasing excitatory inputs would lead to an increase in the number of excitatory synapses onto OPCs, but no change in the number of inhibitory synapses onto OPCs. We found that the number of excitatory inputs onto OPCs increased when neuronal activity was increased, but that the number of inhibitory inputs onto OPCs was unaffected, consistent with our hypothesis. These results indicate both circuit level and synaptic effects of neuronal activity on OPC synapse formation and development.

## DISCUSSION

Knowledge of the role of neuronal activity in regulating synaptic inputs from neurons onto OPCS, as well as whether this is instructive for OPC target selection of neuronal axons and subsequent myelination patterns, is still very incomplete. In order to begin to elucidate this, we aimed to track a localized set of OPCs in different brain regions to understand the extent that neurons contact OPCs in regionally specific ways. We also manipulated neuronal activity to investigate the effect of neuronal activity on OPC responses, specifically examining OPC proliferation, OPC activation, and the types of synaptic proteins that OPCs express when neuronal activity is altered. Finally, we examined whether neuronal synapses onto OPCs are directly altered by changing the tuning of neuronal excitability. We are following up on OPC target selection and subsequent myelination patterns to determine if the neuronal activity is instructive in an additional manuscript in preparation.

Circuits in the cerebral cortex, hippocampus, and striatum are formed by different populations of neurons that integrate input from both local and distant input regions. It is expected that the pattern of myelination will follow a diverse distribution, and is correlated with the activity of each region. Different neuronal subtypes in the cortex exhibit diverse myelination profiles (Xu et al., 2021) (Tomassy et al., 2014). We found that the patterns of these connections were consistent with the expected local circuitry and there were differences in the number of inputs and proportion of inhibitory and excitatory synapses. Similar data has been described before by (Mount & Monje, 2017) in different brain regions such as corpus callosum, premotor cortex, and primary somatosensory cortex. We expanded that analysis to include the dorsal striatum and hippocampus, and it was important for us to investigate which of the systems we would focus our analysis on synaptic contact. We extended the analysis to explore whether there were differences in interneuron inputs by brain region, examining the distribution of neuronal inputs with NeuN, which preferentially marks excitatory synapses, and the distribution of inhibitory inputs with PV and SST, the two major subclasses of inhibitory interneurons in the forebrain. It is intriguing that for all brain regions, neuronal inputs to OPCs followed the patterns of local cytoarchitecture, suggesting that neuron to OPC connectivity likely follows similar programs to build connected networks of cells than neuronal networks. The number of neuron to OPC inputs was a magnitude of order smaller: thousands of granule neurons were connected to each purkinje cell in the cerebellum, and approx. 600 long distance and 20 local neurons were connected to each PV+ IN in the cortex (Wall et al., 2010). This is likely due to a real difference in the number of neuron to OPC connections, but also to differences in viral targeting: unlike the other papers above, we did not use helper viruses to deliver the TVA or G, nor did we use the ROSA-TVA/G mouse, relying instead upon a two-step tTA-dependent activation of a bi-cistronic Tet dependent construct that allowed production of TVA and G in OPCs. We chose this approach to reduce the possibility for recombination in cells other than OPCs, as any neuronal recombination would profoundly alter our results.

Synaptic communication between neurons and OPCs shares common features found in neuron to neuron synapses, such as inhibitory and excitatory postsynaptic receptors, scaffolding proteins, and ion channels expressed in OPCs. Gephyrin and SAP97 are scaffolding proteins from GABAA and AMPA receptors present in OPCs (Chen et al., 2018) (Arellano et al., 2016), and we used these markers to quantify the number of puncta per OPC as a readout of the number of synapses onto OPCs after chemogenetic activation of neurons in the hippocampus. An elevated number of receptors would support the idea that there is a molecular postsynaptic response to the neuronal activation. The question is whether this response is consistent with the sign of neuronal activation: does increased neuronal activation lead to increased excitatory synapses onto OPCs? Or does increased activation result in plasticity in OPC, resulting in a compensatory increase in inhibitory synapses? It appears that the latter may be true: increasing neuronal activity increased Gephyrin+ postsynaptic puncta, characteristic of inhibitory synapses, and decreased SAP97+ postsynaptic puncta, characteristic of excitatory synapses. Surprisingly, we also found that decreasing neuronal activity had similar effects on OPCs: decreasing neuronal activity also resulted in a decrease in the number of SAP97+ synaptic puncta, and an increase in the number of Gephyrin+ synaptic puncta in OPCs. These results are intriguing: our analysis of the effect of chemogenetic inhibition via Hm3Di indicates that the effect on neuronal activity is a clear decrease in the number of spikes in neurons in the hippocampus, and that the opposite is true in Hm4Dq injected animals: there is a clear increase in the number of spikes in the hippocampus (Figure 2). This suggests plasticity in the OPCs as a result of neuronal activity: OPCs may alter the number of inhibitory vs. excitatory synapses to maintain a balance in activation, similar to neurons. We observed that this is the case, at least in animals in which neuronal activity was increased: we found what may be a compensatory mechanism in OPCs, which may be buffering the increased excitation by increasing the number of inhibitory synapses. The composition of excitatory and inhibitory synapses onto an individual OPC is not well understood, nor is it understood how neuronal activity regulates receptor expression or turnover directly. It is clear that plasticity in OPCs is a Ca2+ dependent process (Cheli et al., 2016) and that ion channels and neurotransmitters are highly dynamic in these cells (Spitzer et al., 2019). Future work examining the specific receptor turnover that occurs in response to neuronal activity will be important to expand our understanding of plasticity in OPCs.

To add to this hypothesis, we expected that the proliferation rates would also be increased in this group as described in the literature (Bergles et al., 2010), but we only observed a trend. We are confident that our chemogenetic approach targeted ctip2+ neurons in the hippocampus (Figure 2), so it is unlikely that this is due to an effect of GABAergic-to-OPC synapses, which are known to have no influence on OPC proliferation (Balia et al., 2017). However, increasing AMPARs has been shown not to induce an increase in proliferation (Chen et al 2018) and other works found similar results, with no difference in oligodendrocyte number (Stedehouder et al., 2018). It is still unclear why some neurons, but not others, exhibit activity-regulated myelination, and although our results suggest that the increase in neuronal activation increases the production of receptors, there may be more than one neuronal population that we are capturing with our chemogenetics (neurons that undergo activity dependent myelination and those that don’t. (Almeida et al., 2021) imaged synaptic vesicle fusion in individual neurons in living zebrafish and observed a feedforward model, where they show that axonal vesicular fusion was enriched in hotspots and promoted and consolidated nascent sheath growth. Therefore, the onset of myelination promotes localized axonal vesicular fusion that in turn promoted myelin growth, as well as a higher likelihood for the OPC to respond to specific neuronal synapses. While we did not test the latter in these experiments, our data are consistent with these findings.

Our final hypothesis was that activity dependent neuronal activity would modify neuron to OPC synapses. We hypothesized that there would be an increase in connections between neurons and OPCs when neuronal activity was increased. We did not find this to be the case - there were no differences in the total number of neuron to OPC synapses when neuronal activity was increased or decreased. We found that in 3/4 of our conditions, we had at least one brain in which the total number of neuron to OPC synapses were an order of magnitude higher. It is unclear why this occurred in a subset of animals: all were dosed with the same tamoxifen, were age matched, and received the same dose of DREADD virus and Rabies virus, and counterbalanced for sex. It is possible that there is a small subpopulation of OPCs that are more highly connected to neurons, and that when we sample this heterogeneous population of OPCs, we occasionally target one of these OPCs, resulting in an order of magnitude more OPCs. As this event is rare, it is difficult to postulate what the function of these “super connected” OPCs are. However, even with these super-connectors removed from the sample, we found no differences in the total number of neuronal inputs to OPCs when neuronal activity was increased or decreased. We did however find a change in the subtypes of neuronal inputs that were connected to OPCs. We found that there was no significant change in the number of inhibitory interneurons that are connected to OPCs, either when neuronal activity was increased or decreased. This may have been due to the smaller number of INs connected to OPCs to begin with in the hippocampus; perhaps if we had examined IN inputs to OPCs in the cortex, where there is a larger percentage of IN to OPC connections, we would have found a more robust effect. Importantly, we did find a difference in the number of Ctip2+ excitatory neuron inputs onto OPCs when neuronal activity was increased, and no change when neuronal activity was decreased. This is consistent with the idea that regulating the number of connections onto OPCs is a mechanism for plasticity and for modulating neuronal activity. We are following up on the role of this plasticity in myelination and in myelin node length in a separate study.

The timing of our experiments is likely relevant to the effects that we see in our experiments: our activity dependent manipulations occurred between P28 and P35, in juvenile animals where synapses, circuits, and myelination are still being highly modified, but after the initial peak in developmental synapse formation, OPC proliferation, and myelination. The plasticity mechanisms may look very different at P14, during this initial phase of synapse formation and myelination, or at P7, when OPC proliferation is at its peak. It will be valuable to look at these earlier ages in future studies to be able to test the role of developmental plasticity in neuron to OPC synapse formation and circuit formation.

## Conclusion

In summary, our findings contribute to a better understanding of the mechanisms of OPC development, synapse formation, and circuit formation during juvenile myelination in response to neuronal activation. This study has the potential to direct research toward therapeutic approaches to treat demyelinating diseases, as adaptive myelination and remyelination likely follow similar mechanisms. By manipulating neuronal activity, we discovered the potential to increase myelination in specific regions and different systems, without necessarily depending on cell proliferation. But there is still much to explore: the downstream signaling pathways that are activated in OPCs after neuronal activity modulation, the changes in myelination, both at a single neuron level, and at the level of individual axons, and the role of synaptic activity in regulating these myelination programs. Future work will reveal these mechanisms.

